# Genetic incompatibilities and evolutionary rescue by wild relatives shaped grain amaranth domestication

**DOI:** 10.1101/2023.03.17.533106

**Authors:** José Gonçalves-Dias, Akanksha Singh, Corbinian Graf, Markus G Stetter

## Abstract

Crop domestication and the subsequent expansion of crops have long been thought of as a linear process from a wild ancestor to a domesticate. However, evidence of gene flow from locally adapted wild relatives that provided adaptive alleles into crops has been identified in multiple species. Yet, little is known about the evolutionary consequences of gene flow during domestication and the interaction of gene flow and genetic load in crop populations. We study the pseudo-cereal grain amaranth that has been domesticated three times in different geographic regions of the Americas. We quantify the amount and distribution of gene flow and genetic load along the genome of the three grain amaranth species and their two wild relatives. Our results show ample gene flow between crop species and between crops and their wild relatives. Gene flow from wild relatives decreased genetic load in the three crop species. This suggests that wild relatives could provide evolutionary rescue by replacing deleterious alleles in crops. We assess experimental hybrids between the three crop species and found genetic incompatibilities between one Central American grain amaranth and the other two crop species. These incompatibilities might have created recent reproductive barriers and maintained species integrity today. Together, our results show that gene flow played an important role in the domestication and expansion of grain amaranth, despite genetic species barriers. The domestication of plants was likely not linear and created a genomic mosaic by multiple contributors with varying fitness effects for today’s crops.

## Introduction

Evolution and speciation has long been viewed as a linear process with a single ancestor giving rise to one or more derived species. Genomic data of large samples have revealed ancestry of different species within modern populations in a number of species (Harris and Nielsen 2016; Niu et al. 2019; Chomicki et al. 2020; Kozak et al. 2021; Orlando et al. 2021; Lv et al. 2022). Particularly in plants where reproductive barriers are often weak, the potential for the exchange of genetic material between related species is rather high (Ostevik et al. 2016; Osuna-Mascaró et al. 2023; Sheidai and Koohdar 2023). Yet, the observation of exchanging genetic material between species is often seen as an exception and as a minor contribution to the genomic composition of a species.

Gene flow describes the process of exchanging genetic information between populations or even species (Rieseberg and Burke 2001). While interbreeding between populations might be frequent, the manifestation of gene flow between locally adapted populations or even species is thought to be rare as it would decrease fitness. Nevertheless, beneficial gene flow has been shown to be an important source of variation for local adaptation (Crispo 2008; Sexton et al. 2011; Ellstrand 2014; Tigano and Friesen 2016; López-Goldar and Agrawal 2021). At least partial fertility of hybrids is required, and compatibility between donor and recipient determines the intensity of gene flow (Aguillon et al. 2022). In the course of speciation, the ability to form viable hybrids can be lost, and reproductive barriers that prevent gene flow can evolve. Hence, hybrid incompatibility can hinder gene flow and lead to reproductive isolation. Yet, reproductive isolation in plants is often incomplete or is circumvented by intermediate populations, allowing for gene flow even between different species.

The domestication of crops and animals led to a major transition in human lifestyle and had a profound impact on the genetic makeup of the domesticates (Doebley 2006). Crop domestication can be seen as rapid evolution, often leading to speciation. Even more than speciation in general, crop domestication has long been described as a linear process starting from one wild species evolving through strong directional selection into a domesticate. However, this view has been challenged in recent years, where gene flow from wild relatives have been documented in a number of crops, including maize (Ross-Ibarra et al. 2009), rice (Yang et al. 2012), barley (Civáň et al. 2021), sorghum (Sagnard et al. 2011), tomato (Razifard et al. 2020), *Brassica* (Saban et al. 2023) and others (Luo et al. 2007; Ding et al. 2022; Page et al. 2019; Liu et al. 2019). Reproductive isolation of the crop from its wild relatives would ensure the maintenance of domestication traits, hence, the success of domestication (Dempewolf et al. 2012). Gene flow from wild relatives would have led to a reduction of domestication-related phenotypic changes. Early generations of crop-wild hybrids would be rather unfit as wild plants or crops, as their adaptive traits strongly differ (Janzen et al. 2019; Stetter 2020). Yet, gene flow from wild relatives could have increased the genetic variation in early crops, which could have been beneficial to increase adaptive potential (Smith et al. 2019). In addition, gene flow from locally adapted wild relatives has been shown to have provided alleles that allowed the crop population to establish in novel environments (Van Heerwaarden *et al*. 2011; Hufford et al. 2013; Wang et al. 2021).

Plant domestication has likely been driven by demographic changes and directional selection. Domestication bottlenecks reduced the effective population size, leading to an accumulation of mildly deleterious alleles in the population (Gaut et al. 2018). Selection on major effect domestication genes might have allowed hitchhiking of linked mildly deleterious alleles (Sedivy et al. 2017). Together these effects increased genetic load – the accumulation of deleterious alleles – in the crop population (Bertorelle et al. 2022). Several studies have assessed the accumulation of genetic load in domesticates compared to their wild relatives, e.g., rice (Lu *et al*. 2006; Xu et al. 2006; Nabholz et al. 2014), maize (Rodgers-Melnick et al. 2015; Gaut et al. 2015), and soybean (Kim et al. 2021). While an accumulation of genetic load has been detected in many domesticated species, no increase has been detected in sorghum, potentially due to the transition to selfing in the crop (Lozano et al. 2021). Gene flow from wild relatives with larger effective population size into crop populations might have reduced genetic load in crops (Stetter 2020). In sorghum, gene flow between early landraces resulted in decreased genetic load across landraces, but no variation in genetic load was observed among landraces with or without gene flow (Smith et al. 2019; Lozano et al. 2021). However, such evolutionary rescue by gene flow from wild relatives into domesticates has received little attention. We study the effects of gene flow on genetic load in a three times domesticated crop and its wild relatives to understand the evolutionary role of gene flow and genetic load during crop domestication.

Grain amaranth is a nutritious pseudo-cereal from the Americas that has been domesticated three times from one ancestral species (*A. hybridus*). Two grain amaranths were domesticated in Central America (*A. cruentus, A. hypochondriacus*) and one in South America (*A. caudatus*) (Sauer 1967). The taxonomic complexity of the *Amaranthus* genus led to different domestication scenarios for the crop (Sauer 1967; Kietlinski et al. 2014; Stetter et al. 2017). Population genetic and genome-wide selection signals suggest subpopulations of *A. hybridus* as ancestors for all three grain amaranths (Stetter et al. 2020, Figure 1A and S1). In South America, the closely related *A. quitensis* was potentially involved in the domestication process of *A. caudatus* and genome-wide signals of gene flow between grain amaranths and their wild relatives have been detected previously (Kietlinski et al. 2014; Stetter et al. 2020). While the three grain amaranth species have been cultivated as a crop in different regions of America for thousands of years, all three lack key domestication traits (Stetter et al. 2017). A potential reason for the lack of domestication traits might be continuous gene flow from wild relatives that prevented domestication traits to fix (Stetter 2020). Understanding the underlying genomic signatures of gene flow and selection could improve the understanding of the evolutionary history of crops (Meyer et al. 2012).

**Figure 1.**
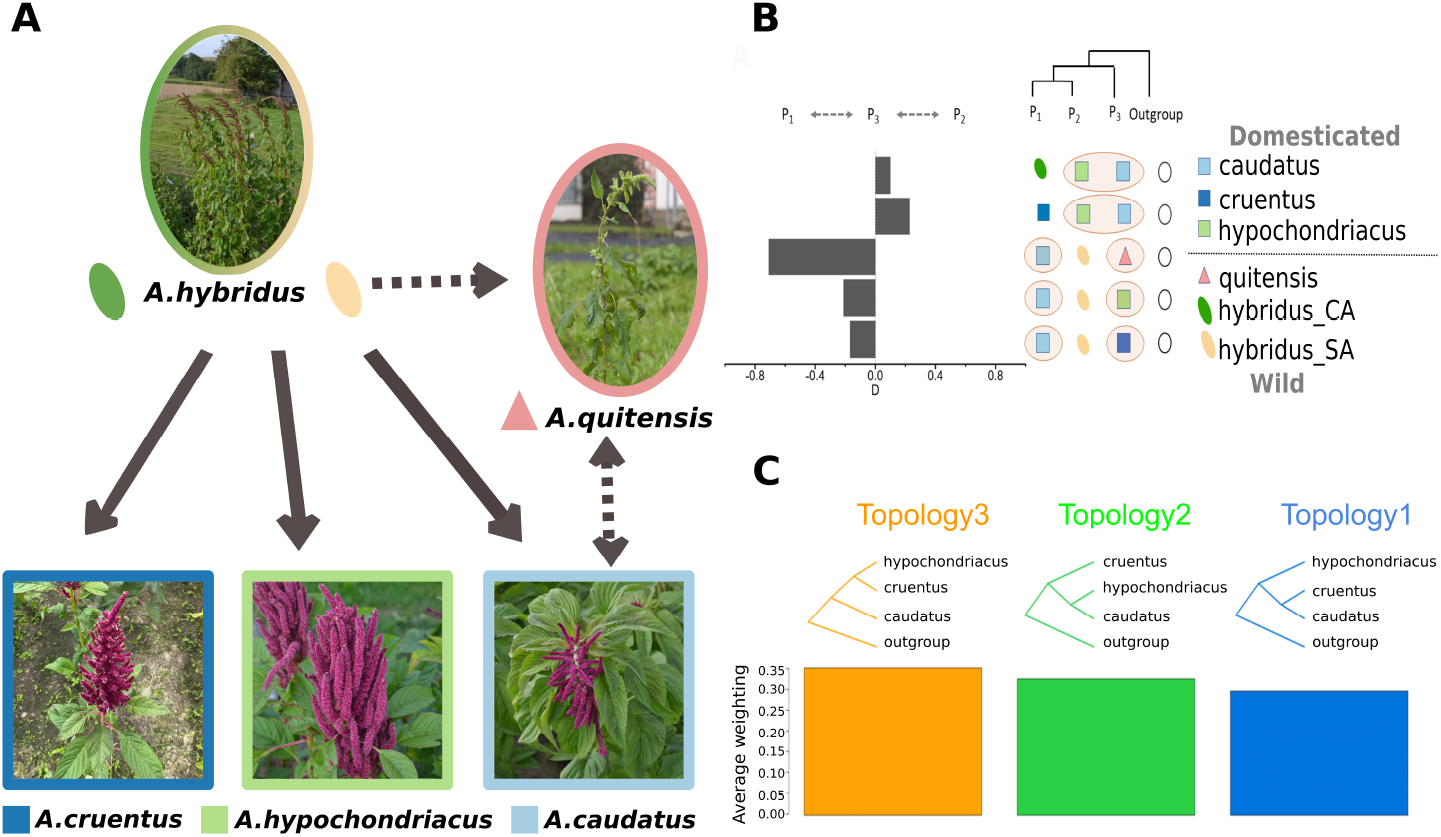
Genome-wide signals of gene flow. A) Schematic history of amaranth domestication. Amaranth has likely been domesticated three times independently from different subpopulations of *A. hybridus* (Central America (hybridus_CA) and South America(hybridus_SA)). *A. quitensis* is speculated to be an intermediate population between *A. hybridus*_SA and *A. caudatus*. Colors are consistent with the legend in B. B) Gene flow between amaranth populations (exchanging pairs highlighted in yellow). The D-value indicates the strength of gene flow. Only significant signals of gene flow are shown. C) Genome-wide summary of tree topologies along the genome inferred by Twisst. The proportion of each of the three topologies observed along the genome is shown in bars.

In this study, we use population-wide whole genome sequencing data and reveal the mosaic signature of gene flow along the genome of domesticated amaranths. We found strong signals of gene flow even between the three geographically isolated domesticated species. Besides post-domestication gene flow between crops, we also observed high levels of genetic exchange between the South American wild species and the local crop. A hybridization experiment between three crops indicates genetic incompatibility between *A. cruentus* and the other two grain amaranths. This reproductive barrier might be a contributing factor to the different rates of gene flow observed between species despite their geographic distances. Gene flow from the wild ancestor *A. hybridus* into the domesticated amaranths reduced genetic load in the crops, but only a few positively selected regions were exchanged through gene flow between crop species. Gene flow might be an important source of genetic variation for crops, not only to provide adaptive alleles but also to reduce genetic load and allow further selection.

## Results

### Strong post-domestication exchange between crops and between crops and their wild relatives

Gene flow likely played an important role during the domestication of different crops (Janzen et al. 2019). In grain amaranth, genome-wide signals of gene flow between species have been previously reported (Stetter et al. 2017, 2020). In order to quantify gene flow among different domesticated and wild populations, we measured ancient admixture using the D-statistic (ABBA-BABA) for all possible tree topologies using ANGSD (Korneliussen *et al*. 2014) in whole genome sequencing data of six population samples of domesticated (caudatus: *A. caudatus*; cruentus: *A. cruentus*; hypochondriacus: *A. hypochondriacus*) and wild amaranth (hybridus_CA: *A. hybridus* from Central America; hybridus_SA: *A. hybridus* from South America and quitensis: *A. quitensis*). We identified gene flow between crop species and between crops and their wild relatives. We found ample gene flow even between geographically distant crop species (Figure 1B). The strongest signal was identified for the Central American crop hypochondriacus and the South American caudatus. This signal was robust even when changing the third species in the test (Figure 1B and Table S1). The test also identified gene flow between the South American crop species caudatus and the second Central American grain amaranth cruentus. However, the strength of gene flow between them was lower. This is also shown when the three grain species were tested in the same tree, where a significant level of gene flow between caudatus and hypochondriacus was identified (D=0.22; Figure 1B, second row), showing that the signal of gene flow between the two species is higher than the shared variation of the three crops. As both Central American grain amaranths were domesticated from Central American hybridus and the exact subpopulations of the ancestor that gave rise to each crop remain unknown, we could not test directly for gene flow between the two Central American grain species. The high level of exchange of genetic material between species was also shown by tree topology tests using Twisst (Martin and Van Belleghem 2017). While the expected tree topology along the genome, with the two Central American crops being closest, was the most common, the other two alternative topologies were only slightly less abundant (Figure 1C). Altogether, more gene flow was observed between two allopatric crop species, caudatus and hypochondriacus, less gene flow was observed with cruentus.

We tested whether isolation with gene flow between hypochondriacus and caudatus was a better fitting model than a simple split without further gene flow by simulating demographic histories using Fastsimcoal2 (Excoffier et al. 2013). We investigated three alternative models: population split without gene flow, population split with one-time gene flow and population split with continuous gene flow (Figure S2). The model for a population split with continuous gene flow was obtained as the best model (Table S2), suggesting that this is the most suitable scenario (Figure S2). In addition, this model predicts population splits estimates that coincide with previous observations (Stetter et al. 2020).

In addition to gene flow between crop species, we examined gene flow signals between the crops and their wild relatives. We found the strongest signal of gene flow between the sympatric South American grain crop caudatus and the local wild quitensis with a D-value of -0.71. When testing a scenario where we assumed quitensis as the ancestor of caudatus, which had been suggested previously (Sauer 1967), significant gene flow from South American hybridus was detected (Figures S3 and S4), indicating the intermediate role of *A. quitensis* between wild and domesticated species.

### Fine-scale gene flow reveals diverse local ancestry of grain amaranths

The genome-wide and population-wide gene flow analysis already showed the complex pattern of exchange of genetic material between the *Amaranthus* species. We found evidence of gene flow for distant and closely related species. To understand the species complex as a whole, we inferred the local ancestry (Lawson *et al*. 2012) along the genome of each individual using finestrucuture v4.1 (Lawson et al. 2012) and summarized them by population (Figure 2). We observed that the Central American grain species cruentus and hypochondriacus had less admixed backgrounds but shared ancestry tracks depending on the different individuals. Both had the highest donated portion from the wild hybridus_CA (Figure 2). Hypochondriacus had a heterogeneous contribution from the other species to different individuals (although with proportions less than 0.016 total), while cruentus had a more homogeneous contribution among individuals in the population. In South America, the overall pattern observed with population-wide introgression tests was also confirmed by the individual-based test (Figures S3 and S4). The South American populations caudatus and quitensis shared large amounts of ancestry, but this strongly varied between the individuals, between 10.8% and 18.5% of quitensis ancestry in caudatus individuals(Figure 2A), which also varied along the genome (Figure 2B).

**Figure 2.**
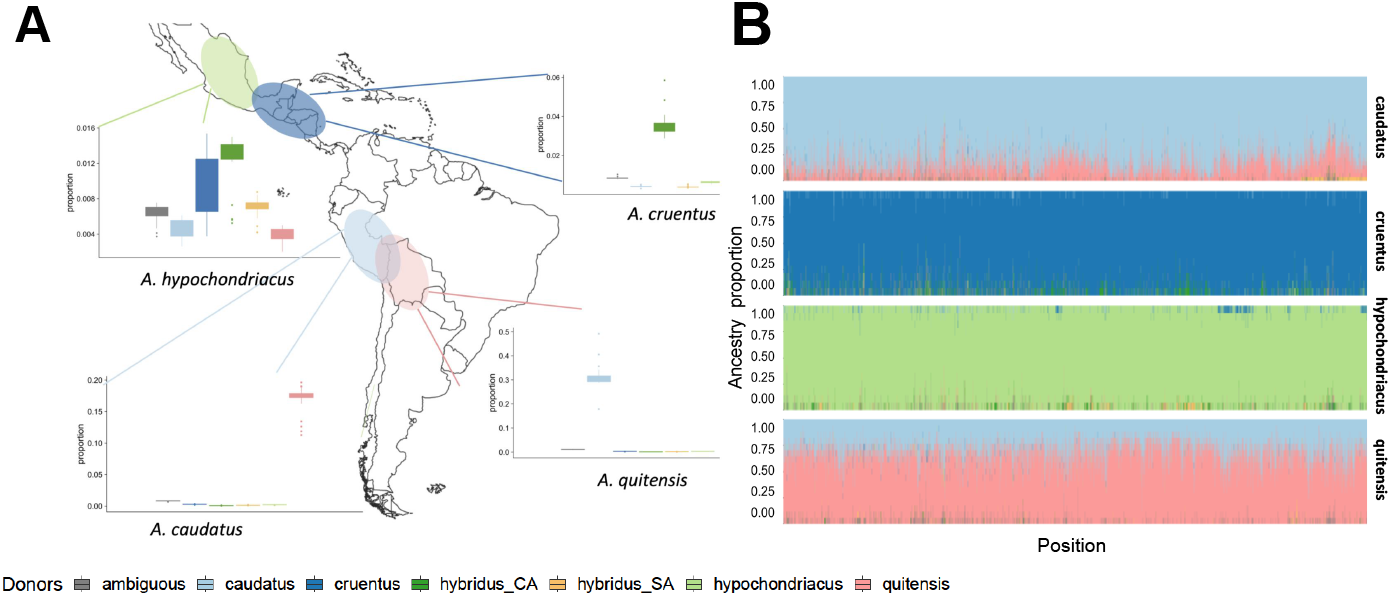
Variable ancestry across individuals and along the genome. A) Contribution of donor populations to individual recipients. Values within boxplots represent contributions by different donor populations to each individual of the recipient population. The Y-axis scale differs between plots. The schematic geographic range of populations. B) Population scale ancestry proportion along genomic positions of Scaffold 4. The proportion of the most likely donor population at a given SNP across all individuals in the recipient population. Each plot represents a recipient population, and colors represent donor populations. Exemplary scaffold, all scaffolds in Figure S5. Donor colors according to Figure 1.

We further wanted to understand whether gene flow is variable along the genome. Comparing ancestry signals along the genome of the different species showed that even the same region can have multiple donors in a species (Figure 2B and Figure S5). Using D-value in genomic windows, we scanned the genomes for gene flow signals in the trees with significant genome-wide signals. The previous comparison between crops showed the presence of gene flow between hypochondriacus and caudatus and between cruentus and caudatus (Figure 1). In the local scan, we observed similar signals as for global gene flow analysis; stronger gene flow between caudatus and hypochondriacus (D>0.992 for top 1% windows) than between caudatus and cruentus (D < -0.887 for bottom 1% windows) (Figure 3). Windows representing significant gene flow from cruentus or hypochondriacus with caudatus did not overlap, suggesting that gene flow occurred independently between species.

**Figure 3.**
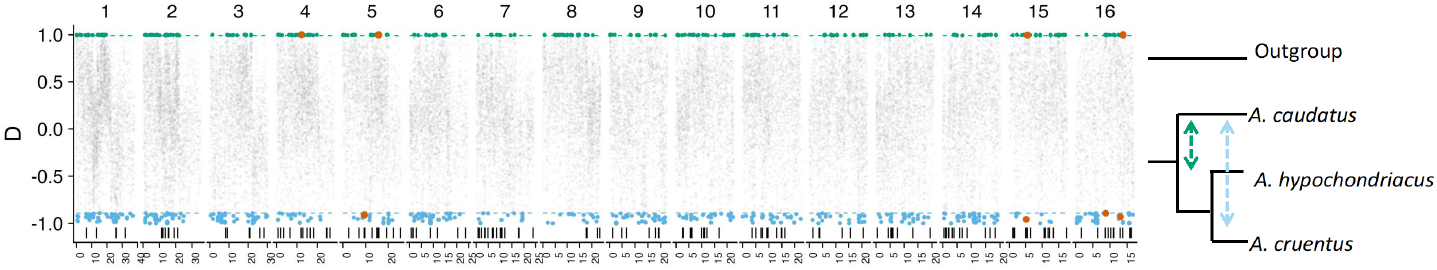
Fine-scale gene flow along the genome between domesticated amaranth populations. We used the D-value to calculate gene flow along the genome in 1000 SNPs windows. Each dot represents the D-value for a tree comparing *A. caudatus* with *A. hypochondriacus* or *A. cruentus*. Positive values are indicative of gene flow between *A. caudatus* and *A. hypochondriacus* and negative values between *A. caudatus* and *A. cruentus*. The top 0.1% of windows in each direction were colored. The bars at the bottom indicate previously detected selective sweep regions detected in *A. caudatus*. Orange dots denote overlaps between selective sweep and top gene flow signal.

To understand the potential reason for the observed high levels of gene flow between grain amaranth species, we combined gene flow signals along the genome with selection scan results (Gonçalves-Dias and Stetter 2021). Despite the genome-wide distribution of gene flow between crop species, only a few selective sweeps overlapped with outlier windows of gene flow between caudatus and the other two crop species. We found 13 overlapping windows in total, eight in regions of gene flow between hypochondriacus and caudatus and five between caudatus and cruentus. Despite the relatively low total number, the overlap was higher than expected by chance (p=0.02), suggesting beneficial gene flow between geographically distant crop relatives.

### Introgression from wild ancestor mitigates increased genetic load in domesticated grain amaranths

In many crop species, a reduction in overall genetic diversity between wild relatives and the crop has been observed. This has been associated with population bottlenecks and directional selection during domestication (Gaut et al. 2018). Increased genetic drift and hitchhiking with selected alleles can lead to a higher genetic load in the domesticated species (Lu et al. 2006; Wang et al. 2017). We calculated GERP scores for the *A. hypochondriacus* reference genome from whole genome alignments with 15 diverse plant species of different relatedness as a proxy for deleterious alleles (Figure S6). We observed that two domesticated species (caudatus and cruentus) had a significantly higher total genetic load than their wild ancestor hybridus. The third crop species, hypochondriacus, had a higher load than hybridus_SA but lower than hybridus_CA (Figure 4A). This pattern remained even when using the *A. cruentus* reference genome (Ma et al. 2021) to calculate GERP scores, showing that the difference is likely not the result of reference bias when calculating GERP scores (Figure S7). The South American relative quitensis showed as high total load as the domesticated species. Partitioning of total genetic load from fixed and segregating sites showed that the domesticated species had a high fixed load but a lower segregating load than their wild ancestor (Figure 4B and C). Quitensis also showed high fixed load and low segregating load, in agreement with the small effective population size and low genetic diversity documented previously (Stetter et al. 2020).

**Figure 4.**
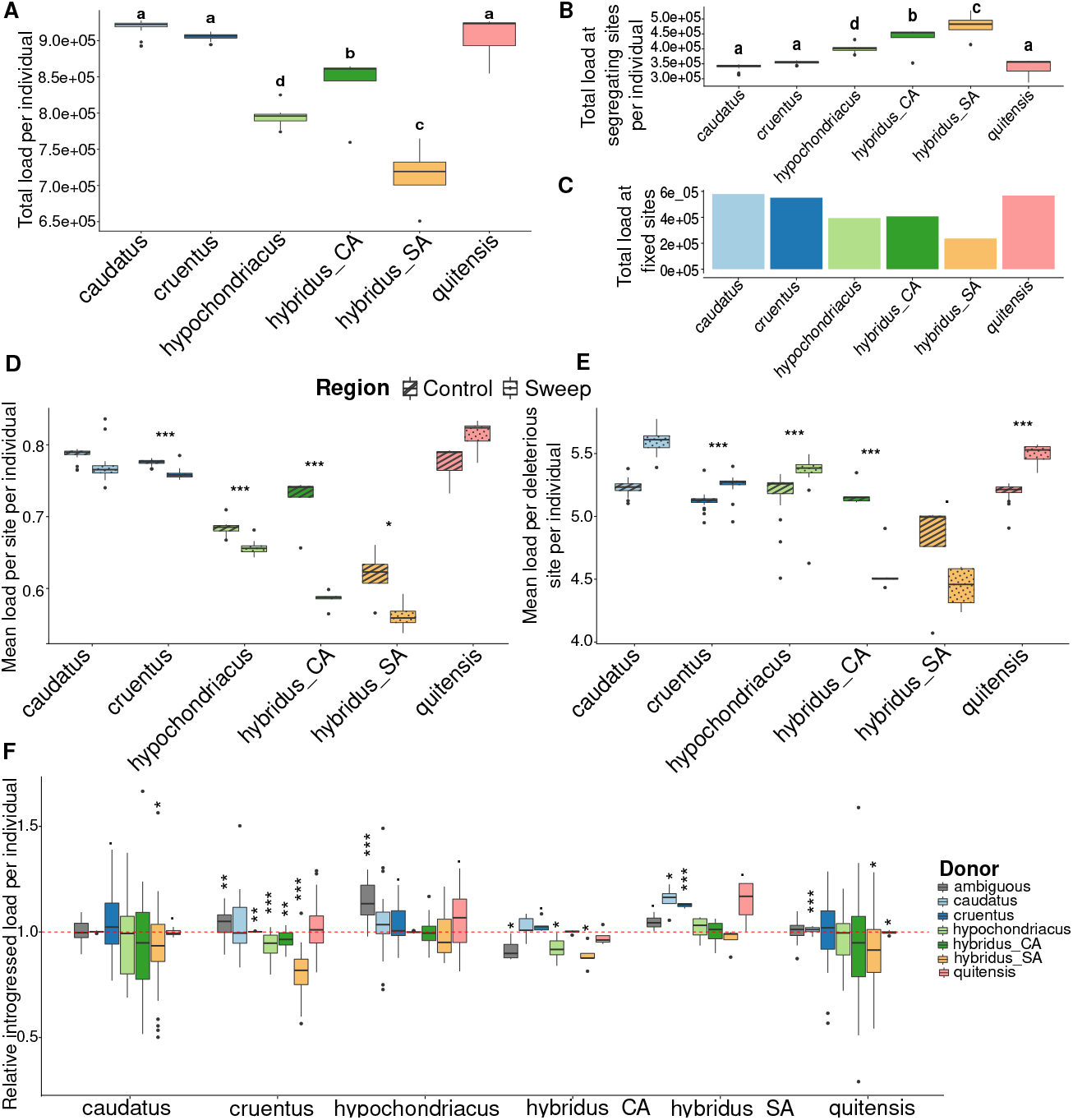
Genetic load in domesticated and wild amaranth. A-C) Genetic load was calculated as the sum of GERP scores for the derived allele per individual. A) Total genetic load per individual in domesticated and wild populations. Different letters above the box plot represent significant differences. B) Segregating genetic load in each population per individual and C) Fixed genetic load within each population. D) and E) Accumulation of genetic load in selective sweep regions and control regions (rest of the genome). D) Mean load of sweep/non-sweep region (including all sites in region); E) Mean effect (GERP score) of deleterious allele in region (only deleterious sites). Asterisks above box plot represent the significance level. F) Relative introgressed genetic load; load in introgressed regions relative to introgression received from the donor. A value greater than one represents increased load through introgression, while a value lower than one shows a reduction in load through introgression compared to the amount of introgression by the donor. A value of 1 shows, the expectation of equal load and introgression proportion (denoted by red dotted line). The asterisks above the box plot represent the significance level for one-sample t-test. (* - p-value < 0.05, ** - Pvalue < 0.01, *** - Pvalue < 0.001)

To investigate if hitchhiking of deleterious alleles with selected loci led to the overall increase in genetic load in the domesticated species, we compared genetic load within selective sweep regions (Gonçalves-Dias and Stetter 2021) with that of random non-sweep regions. We observed that the mean load per site in sweep regions was significantly lower than in control regions for all domesticates (Figure 4D). However, the mean GERP score per deleterious site in the sweep regions showed higher values than deleterious sites in control regions, suggesting a hitchhiking effect (Figure 4E). This suggests that strongly deleterious alleles might accumulate within selective sweep regions due to hitchhiking, but mildly deleterious alleles fix or increase in frequency due to increased genetic drift.

The abundant gene flow between grain amaranths and their wild relatives is expected to impact the patterns of genetic load. To evaluate the potential effect of gene flow on the accumulation of deleterious alleles, we measured the total load accumulated within introgressed regions from other species into a recipient species as the proportion of introgressed load received per individual. The expected value if introgressed regions carry the same amount of load as random regions would be one, while the value is higher than one for deleterious introgression and less than one for beneficial introgression. The analysis showed that introgressed regions from *A. hybridus* into the domesticated reduced genetic load, while introgression between crops donated higher load in the recipient population (Figure 4F). Introgression from domesticated donors into the wild species resulted in a higher genetic load. These results might suggest that gene flow from populations with higher effective population sizes (wild relatives) could provide evolutionary rescue for the smaller populations (crops).

### Hybrid incompatibilities between grain amaranths

All gene flow begins with the inter-mating between individuals of different populations. To further understand the process of gene flow and differences in observed levels of gene flow, we crossed multiple inbred lines of the three crop species. All three amaranth species are mostly selfing, but outcrossing is possible and occurs in the field. We selected three accessions from each crop species and crossed them within and between species, including selfings. All crosses produced viable F_1_ seeds. Selfings and intra-specific F_1_ plants grew without complications and set seeds (Figure 5, Table S3). The inter-specific F_1_ plants of the 5 combinations between *A. caudatus* and *A. hypochondriacus* developed healthy and fertile plants. For crosses between *A. cruentus* and *A. hypochondriacus*, one of the three combinations led to a lethal phenotype, with unhealthy seedlings that did not survive the juvenile stage (Figure 5). Yet, both parents of this cross produced healthy and fertile offspring with other crossing partners, suggesting that incompatibility is the result of a specific allelic combination rather than an individual unfit parent (Table S3). All of the six combinations between *A. caudatus* and *A. cruentus* resulted in the lethal phenotype, suggesting a genetic incompatibility between these species. The difference in hybrid compatibility between grain amaranth species likely contributed to the observed patterns of gene flow.

**Figure 5.**
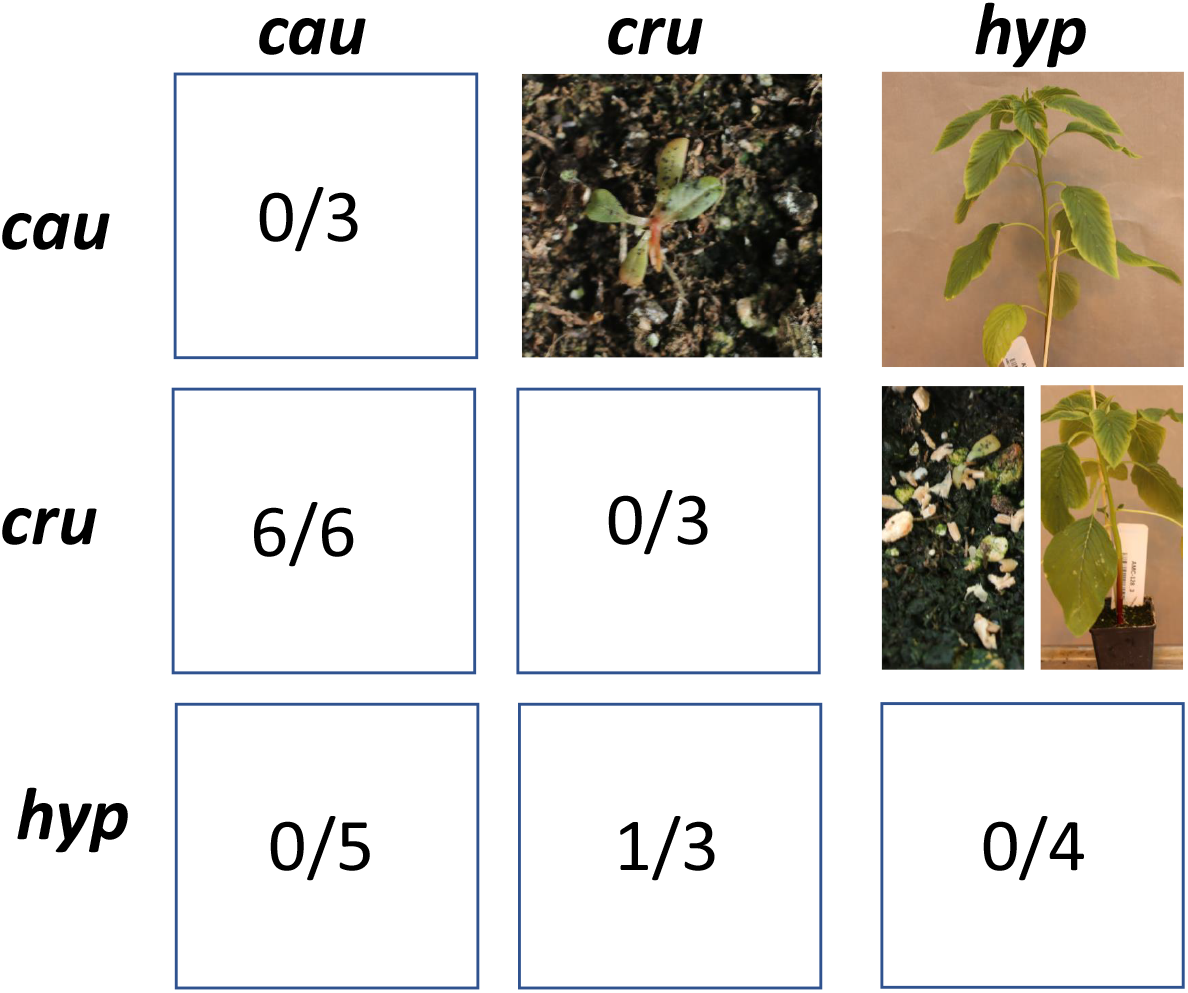
Proportion of lethal F_1_ phenotypes. The lower triangle shows the number of lethal combinations between accessions out of the total number of combinations. We considered a cross as “lethal” when all F_1_ seedlings died within 20 days after planting. The upper triangle shows example images of phenotypes of inter-specific combinations. cau: *A. caudatus*, cru: *A. cruentus*, hyp: *A. hypochondriacus*

## Discussion

Crop populations that we observe today are the result of different evolutionary processes. While selection and demographic changes have been extensively studied, gene flow and its role in the fitness of crops has only received attention in recent years. This is partially due to technical advances in plant genomics, but might also have resulted from the conceptual assumption of a linear process from one wild ancestor. The complex makeup of modern grain amaranth shows that gene flow between crop populations, even over long geographic distances, was prevalent (Figure 1). Gene flow between crop lineages of species that were domesticated multiple times has also contributed to diversity in rice (Yang et al. 2012), tomato (Razifard *et al*. 2020) and common bean (Rendón-Anaya et al. 2017). Not only does such gene flow between closely related crop populations occur, it is also heterogeneous between individuals and along the genome (Figure 2).

A potential reason for lower genetic exchange between specific pairs of grain amaranth could have been the reported difference in chromosome number between *A. cruentus* with 17 chromosomes in comparison to 16 chromosomes in the other 4 species (EJ and Poggio 1994; Ma et al. 2021). Yet, this would only lead to infertile F_1_ plants, rather than necrotic, non-viable plants that die in the seedling stage. Instead, *A. cruentus* formed fertile hybrids with two out of three *A. hypochondriacus* accessions, suggesting incompatibility not because of difference in chromosome numbers but rather a genetic incompatibility (Figure 5). Our crossing experiment revealed differential genetic incompatibility between grain amaranth species, consistent with previous observations on interspecific hybrid necrosis (Gupta and Gudu 1991). A potential one-locus underdominance model of hybrid incompatibility would require strong genetic drift in both populations (Wu and Ting 2004), which could be the result of previously demonstrated domestication bottlenecks in grain amaranth (Stetter et al. 2020). The observed reproductive barrier could also be the result of a Dobzhansky-Muller incompatibility (Muller 1942), including more than one locus. Given the large geographic distance between incompatible crop species (*A. caudatus* in South America and *A. cruentus* in Mesoamerica) the reproductive barriers likely evolved through neutral processes rather than selection against gene flow. The incomplete barrier between *A. cruentus* and *A. hypochondriacus* might allow further insights into the progression of reporductive isolation during crop domesticaiton (Tenaillon et al. 2023). The genetic mechanism for incompatibility warrants further investigation, as this has practical implications for potential hybrid breeding using different crop species as heterotic pools. The complete compatibility between *A. hypochondriacus* and *A. caudatus* is reflected in higher gene flow signals than between these species and *A. cruentus* (Figure 1 and 5). Therefore, *A. hypochondriacus* and *A. caudatus* would be the most promising heterotic pools for future amaranth breeding.

Given the high prevalence of incompatibility between *A. cruentus* and *A. caudatus*, gene flow might have occurred early during the 8,000 year-long domestication history if strong reproductive barriers only developed later in the process. If the hybrid incompatibility arose at the advent of amaranth domestication, it can be expected that gene flow between *A. cruentus* and *A. caudatus* was likely beneficial, given the high fitness disadvantage of F_1_ hybrids (Janzen et al. 2019; Aguillon et al. 2022). We found a low but significant number of introgressed selective sweeps that might represent such beneficial gene flow between amaranth crop species. Currently, there are only a few domestication-related QTL known in grain amaranth that would allow linking introgressed regions to phenotypic changes during domestication. The previously reported QTL for the seed color change during amaranth domestication did not show signals of introgression consistent with the previously reported repeated selection for the trait in the three crop species (Stetter et al. 2020). More quantitative genetic and functional analyses could reveal additional QTLs that could indicate the adaptive potential of introgressed regions between grain amaranths.

Adaptive gene flow from wild relatives into crops has previously been associated with environmental adaptation in crops. For instance, in maize, the introgression of the wild relative *Zea mays spp. mexicana* has been associated with the adaptation of maize to highland conditions and colder climates (Wang et al. 2017). Recent work even suggests a prevalent role of *Zea mays spp. mexicana* in the domestication of maize (Yang *et al*. 2023). While we cannot associate gene flow from wild relatives with positive selection, we found decreased genetic load in regions that were introgressed from wild relatives (Figure 4). This could be due to higher effective population size and higher genetic diversity of wild relatives (Stetter et al. 2020). The wild ancestor *A. hybridus* showed lower genetic load than the domesticates (Figure 4), which might be the result of less demographic change during the recent past (Stetter et al. 2020). Post-domestication gene flow between crops and their wild ancestor could consequently reduce the frequency of deleterious alleles. A similar correlation of genetic load and gene flow from a wild relative has also been shown in maize and sunflower, where gene flow regions from the wild relative showed reduced genetic load (Wang et al. 2017; Huang et al. 2023). Similarly, work in humans has shown that gene flow from a relative with a small population size (Neanderthal) into a population with a larger population size (modern humans) led to increased genetic load (Harris and Nielsen 2016), as we observe for gene flow from domesticated amaranths into *A. hybridus*. The accumulation of genetic load in populations with small effective population sizes can even lead to the extinction of populations or species as a whole (Rogers and Slatkin 2017). Hence, gene flow from relatives with large population size not only provides adaptive variation but can also lead to the evolutionary rescue of the small population (Carlson et al. 2014). Despite the amelioration of genetic load through gene flow with wild relatives, crop-wild hybrids are expected to perform poorly as crops. Gene flow, therefore, needs to have an overall beneficial effect that is higher than the destruction of domestication traits (Stetter 2020). This might be particularly possible in crops like grain amaranth where the domestication syndrome is only weakly pronounced (Stetter et al. 2020).

For crops that maintain high gene flow with their relatives and are phenotypically less differentiated from wild plants, as is the case for grain amaranth, the borders between wild and domesticate might be fluid. Strong gene flow between wild and domesticated crops might also allow the domesticate to return to the wild and be viable without human intervention again. Such feralization has been observed for a number of plant and animal species (Gering et al. 2019). We found particularly strong signals of gene flow between grain amaranth and its wild relatives in South America (Figure 1B). The close relationship between wild species and crop in South America has led to different hypotheses for the domestication of *A. caudatus* suggesting *A. quitensis* as potential wild ancestor (Sauer 1967). While this cannot be completely ruled out, previous work using genome-wide makers data suggested *A. hybridus* as ancestor for all three grain amaranths (Kietlinski et al. 2014; Stetter et al. 2020). The high and genome-wide equally distributed signal of gene flow between *A. caudatus* and *A. quitensis* (Figures 1 and 2) together with the low population size and signs of a strong population bottleneck in *A. quitensis* (Stetter et al. 2020) might indicate a feralized status of this species. The clarification of the status of *A. quitensis* will need further work with multiple populations of this species, local crop and wild relatives.

Overall, we show that the relationships between species are beyond linear, with exchanges between populations despite large geographic distances and the reintroduction of genetic material that was potentially lost during speciation. Even with observed genetic incompatibilities and high genetic differentiation between the crop species, we found strong signals of gene flow between grain amaranths and between the crops and their wild relatives. Recurrent gene flow from the wild relative into the crops might have allowed evolutionary rescue, counteracting the loss of diversity, but likely hindered the fixation of domestication traits leading to the incomplete domestication syndrome observed today for grain amaranth.

## Materials and Methods

We studied whole genome resequencing data of 108 domesticated and wild amaranth accessions. The raw reads are available from European Nucleotide Archive (project numbers PRJEB30531) (Stetter et al. 2020). The accessions included the three domesticated amaranth; 33 *A. caudatus* L. (caudatus), 21 *A. cruentus* L. (cruentus) and 21 *A. hypochondriacus* L. (hypochondriacus); as well as 5 wild *A. hybridus* L. from Central America (hybridus_CA), 4 *A. hybridus* L. from South America (hybridus_SA), and 24 *A. quitensis* Kunth. (quitensis) (Table S5). *A. tuberculatus* was used as outgroup for the study (ERR3220318), (Kreiner et al. 2019). Raw reads were aligned to amaranth reference genome (Lightfoot et al. 2017) using bwa-mem2 (v 2.2.1) (Vasimuddin *et al*. 2019).

### Variant calling

For variant calling, we utilized ANGSD (v.0921) (Korneliussen et al. 2014), with -ref *A. hypochondriacus* V2.1 reference genome (Lightfoot et al. 2017) - doCounts 1, doGeno 3 dovcf 1, gl 2, dopost 2, domajorminor 1 anddomaf 1. We filtered for missing data and mapping quality using minInd 73 (max 30 missing data), minQ 20, minMapQ 30, only_proper_pairs 1, trim 0, SNP_pval 1e-6, setMaxDepthInd 150 and setminDepth 73. The resulting VCF file was phased using Beagle (v 5.2) (Browning et al. 2021) using default parameters. For linkage disequilibrium (LD) pruning, we used Plink (v 1.9) (Purcell et al. 2007) using windows of 50kb with 5kb steps and a r^2^ threshold of 0.3. The resulting VCF file had a total of 13,330,082 sites.

### Gene flow analysis

We inferred gene flow between populations using D-statistic implemented in ANGSD (v.0921) (Korneliussen et al. 2014). We inferred population-wide statistics with the abbababba function for calculations of D per individual and abbababba2 for calculations between populations. For both tests, *A. tuberculatus* was used as outgroup (H4). Only trees with a significant Z-score (absolute value above 3) were included in the results. For fine-scale analysis along the genome, we employed Dsuite (Malinsky et al. 2021-02). We used the function Dinvestigate in windows of 100 SNPs to calculate D between trios along the genome. We also utilized the function Dtrios to verify the concordance of the global genome with the results obtained from ANGSD. To overlap regions with gene flow between crop species with selection signals, windows with significant signals of gene flow (1% outlier values) were overlapped with selective sweeps signals in the recipient population. Selective sweeps were previously identified in Gonçalves-Dias and Stetter (2021). The overlaps were tested for significance using a hypergeometric test (*pyhper* function in R 4.2).

### Topology inference

We used Twisst (Martin and Van Belleghem 2017) to infer the topology of each trio along the genome in windows of 100 SNPs, utilizing *A. tuberculatus* as an outgroup. For each window, a topology is assigned and a summary of the proportional windows in which each topology appeared is then obtained. This inference allows a blind observation of the relationship between species. In the case, where topologies that differ from a neutral expectation are present in high proportions suggests gene flow between species. We inferred the topology for trios, between which a putative gene flow signal was identified.

### Local Ancestry inference

We inferred local ancestry for each individual using finestrucuture v4.1 (Lawson et al. 2012). We used a uniform recombination rate generated with the perl script makeuniformrecfile.pl. The program was run using parameter -f 0 0 which considers the populations and iterates through all individuals. For each individual, all five species (including individuals of the same species) were used as donors, which allowed to differentiate ancestry from all the species. We further assigned the most likely donor for each genomic region of an individual. The donor for each position per individual was assigned based on the most likely donor population with a likelihood larger than 0.5. Sites that could not meet these likelihood thresholds were called “ambiguous”. Using these thresholds, the proportion of donated region per individual was calculated. In addition, the proportion along the genome within each population was summarized.

### Demographic modeling

To estimate whether the identified scenario of gene flow fits best to our data we used simulations using Fastsimcoal2 (Excoffier et al. 2013). The joint site frequency spectrum (SFS) was generated using non-coding SNPs having no missing value in any of the individuals and a minimum coverage of five reads using a python program easySFS (https://github.com/isaacovercast/easySFS). First, the program was run on preview mode (–preview) to identify the true sample size and the best sample size selected was used for the projection (–proj) to generate the joint SFS. Three different models namely, two-population split, two-population split with one migration event and two-population split with continuous gene flow (Figure S2) were applied to the observed joint SFS. The models were compared using Akaike’s Information Criterion (AIC). The best parameter estimate was calculated based on 100 independent runs with 200,000 coalescent simulations and 40 cycles of likelihood maximization algorithm. The 95 percent confidence interval of each parameter was estimated based on 50 non-parametric bootstrapping datasets. Each of the 50 bootstrapped datasets was run 50 times to estimate the best run. These 50 best run parameters were then used to estimate the confidence interval using the boot package in R (Canty and Ripley 2017).

### Genetic Load

We used Genomic Evolutionary Rate Profiling (GERP) (Davydov et al. 2010) scores to account for the effect of deleterious alleles at each site. We aligned 15 repeat-masked genomes of angiosperm species spanning a large taxonomic range to the reference genome of *A. hypochondriacus* (v2.1) (Lightfoot et al. 2017). We followed the pipeline of Wu *et al*. (2022). Briefly, we aligned genomes of 16 divergent species: *Beta vulgaris* EL10 1.0, *Brachypodium distachyon* v2.1, *Chenopodium quinoa* v1.0, *Glycine max* Wm82.a4.v1, *Helianthus annuus* r1.2, *Medicago truncatula* Mt4.0v1, *Mimulus guttatus* v2.0, *Oryza sativa* v7.0, *Phaseolus vulgaris* v2.1, *Populus trichocarpa* v4.1, *Setaria viridis* v2.1, *Solanum lycopersicum* ITAG4.0, *Sorghum bicolor* v3.1.1, *Spinacia oleracea* (Monoe Viroflay) and *Vitis vinifera* v2.1 from phytozome (Goodstein et al. 2012) to the *A. hypochondriacus* reference genome using the LAST aligner (Kiełbasa et al. 2011). The tree topology for the species was extracted from the NCBI-phylogeny using the ete3 toolkit (v3.1.2) (Huerta-Cepas et al. 2016). Phylofit from phast package (Siepel et al. 2005) was used to calculate the branch length of the tree along with four-fold degenerate sites from *A. hypochondriacus* reference annotation file to generate a neutral model. All pairwise alignments were merged using ROAST (https://github.com/multiz/multiz) (Blanchette et al. 2004). GERP++ (Davydov et al. 2010) was used to calculate the GERP scores for each site using gerpcol with -j option that projects out the reference genome to avoid bias in the calculation. All sites with negative GERP values were set to 0 as negative values are not informative and misleading. The ancestral allele for each site was defined on the basis of three outgroup species closest to *Amaranthus* in the phylogenetic tree, i.e., *Beta vulgaris, Chenopodium quinoa* and *Spinacia oleracea*. Variants among these three species were called from the roast multiple alignment file (maf) using maffilter (version 1.3.1) (Dutheil et al. 2014). Sites not covered by any of the three outgroup species were removed. For the remaining sites, the major allele among the three species was called the ancestral allele. In case of discrepancy for a majority rule, the allele for *Spinacia oleracea* was called as the ancestral allele.

From a total of 13.2 million SNPs, we extracted 1,429,744 sites for which the GERP score and the ancestral alleles could be assigned. We then polarised the SNPs and removed sites where neither the reference nor the alternate allele matched the ancestral one. This yielded a total of 1,145,566 SNPs that were used for genetic load calculation. The GERP score from sites having derived alleles was then summed to calculate the total genetic load using an additive model. The genetic load was calculated for each accession individually. The significance of variation for the load in each population was then analyzed using an ANOVA followed by TukeyHSD in R (https://www.rdocumentation.org/packages/stats/versions/3.6.2/topics/TukeyHSD).

To account for the reference bias, the GERP score was also calculated using *A. cruentus* genome (Ma *et al*. 2021) as reference, using the same method as described above. The variants file of individuals that used *A. hypochondriacus* as reference was lifted to the *A. cruentus* genome using liftover module of maffilter (version 1.3.1) (Dutheil et al. 2014). The individual genetic load was then calculated using GERP score from *A. cruentus* genome for each lifted site as described above.

To overlap genetic load and selective sweeps, the sweep regions for each of the populations were overlapped with the sites with GERP score. Variants from those regions were extracted for the individuals of the respective populations. Any other regions of the genome that were not under the sweep region were considered as control region. To account for the difference in the sizes of the sweep and control regions, we divided the total load by the total number of sites used in the analysis. Differences in genetic load between sweep and non-sweep regions were then tested using a t-test for each population.

In order to calculate the introgressed load, GERP scores were summed for sites having the derived allele donated by a different population. The introgressed load was expressed as a ratio of load contributed by the introgressed species to the percentage of introgression sites. A value greater than one predicts the contribution of a higher load due to gene flow. The significance of deviation from the expected value was analyzed using one-sample t-test against the null-expectation of equal contribution of 1.

### Experimental hybridization between grain amaranth species

We selected genetically and morphologically defined accessions for each of the three crop species (*A. caudatus, A. cruentus*, and *A. hypochondriacus*) to assess their cross-compatibility. We crossed 9 parental accessions (three per species) to create 25 inter - and intraspecific combinations and examined multiple crosses per combination (Table S3). The parental lines were previously selfed for at least three generations to ensure homozygosity. As the three grain amaranths are mostly selfing, we hand-emasculated the female parent and bagged parents together. Successful crossing was ensured by PCR using diverging primer pairs (Table S4). To determine the hybrid survival rate, the hybrids were grown alongside their parental accessions in a greenhouse in Cologne (Germany) under long day conditions (16h light, 8h dark) at 25°C. We evaluated the survival of hybrid plants from at least three offsprings per cross. We considered a cross as “lethal” when all F_1_ seedlings died within 20 days after planting.

## Data Availability

Genomic data is available through Stetter *et al*. (2020) and the associated ENA project. All scripts used in the analysis are available on https://github.com/cropevolution/GeneFlowLoadRescue.

## Author contribution

MGS conceived the study. JGD processed the data and conducted genome-wide analysis of gene flow and local ancestry inferences. AS performed genetic load analysis and demographic modeling. CG performed experimental crosses and executed the incompatibility study together with JGD. JGD, AS and CG prepared figures and tables. MGS, JGD and AS wrote the manuscript. All authors discussed the results, edited and approved the manuscript.

## Acknowledgments

We acknowledge funding by the Deutsche Forschungsgemeinschaft (DFG, German Research Foundation) under Germany’
ss Excellence Strategy – EXC-2048/1 – Project ID 390686111 and grant STE 2654/5 to MGS by the DFG.

