## Supplementary material for "Genetic incompatibilities and evolutionary rescue by wild relatives shaped grain amaranth domestication": S1

### Supplementary Figures

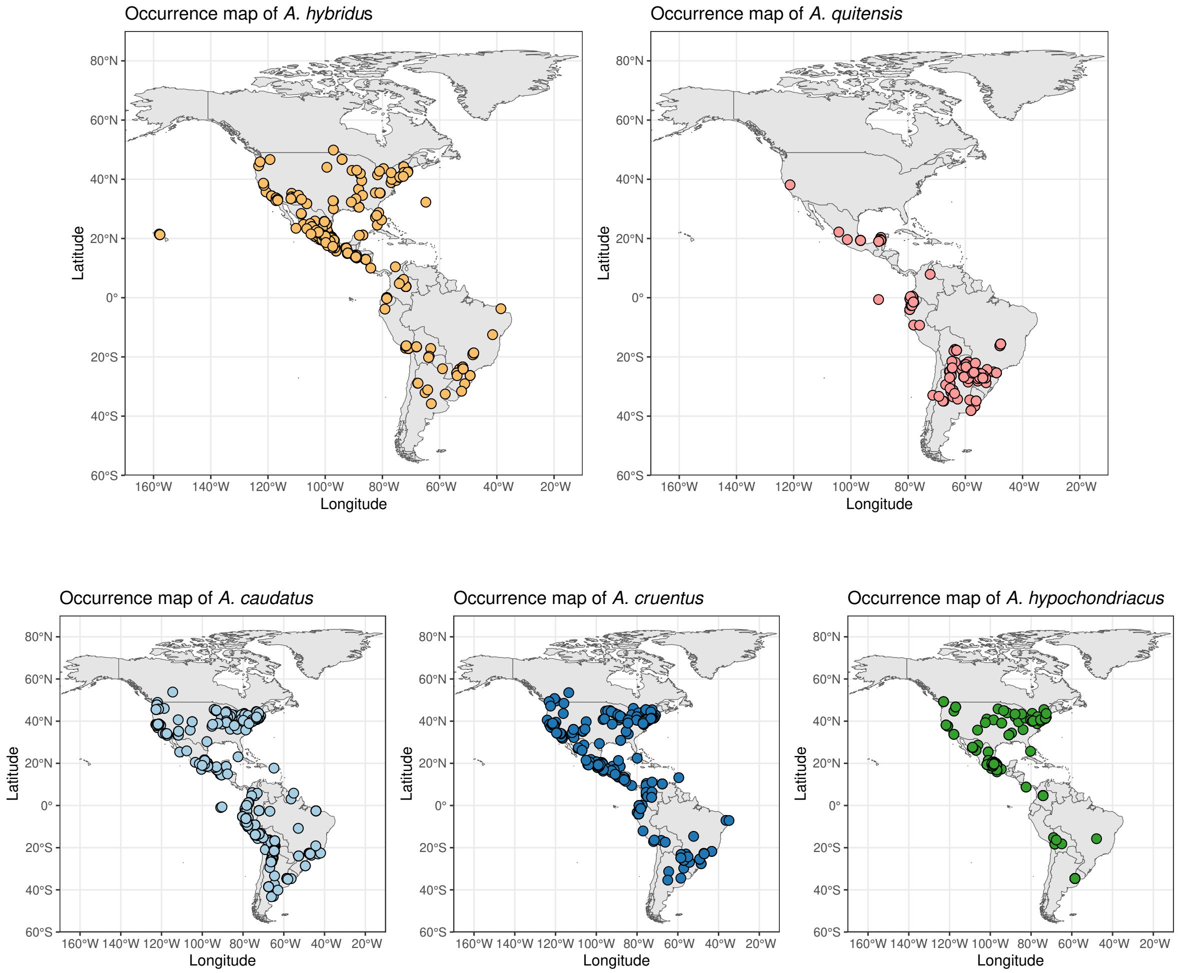


Figure S1: Occurrence map for two wild and three domesticated species of grain amaranth. For *A. hybridus*, the samples in the Central American and South American regions were denoted as hybridus_CA and hybriuds_SA, respectively with no clear range distinction. The data for occurrence were extracted from the GBIF database (https://doi.org/10.15468/dd.f8cz3g).

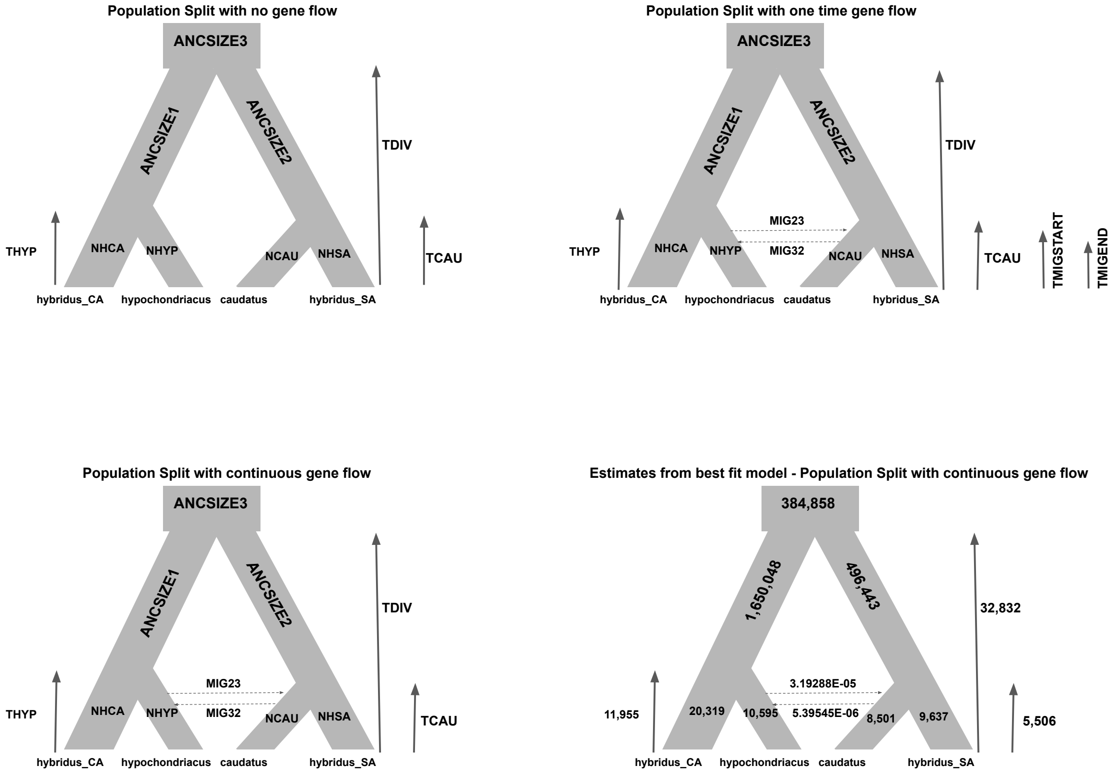


Figure S2: Different demographic models used in Fastsimcoal2 to predict the best scenario. The model of population split with continuous gene flow was predicted to be the best model. Also see Table S2 for detailed demographic parameters for each model.

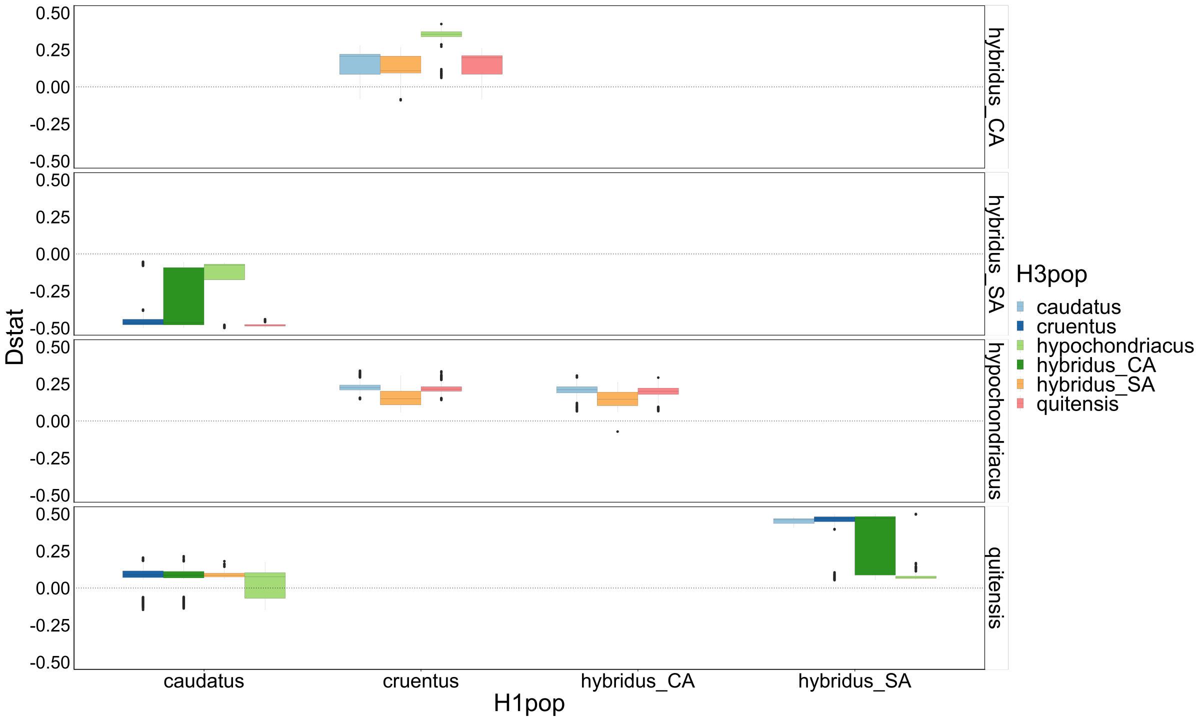


Figure S3: D-statistic value for comparisons between individuals. Each boxplot represents a comparison for a population trios H3 with the inner node H1 and H2. Only significant trees are represented. *A. tuberculatus* was used as an outgroup.

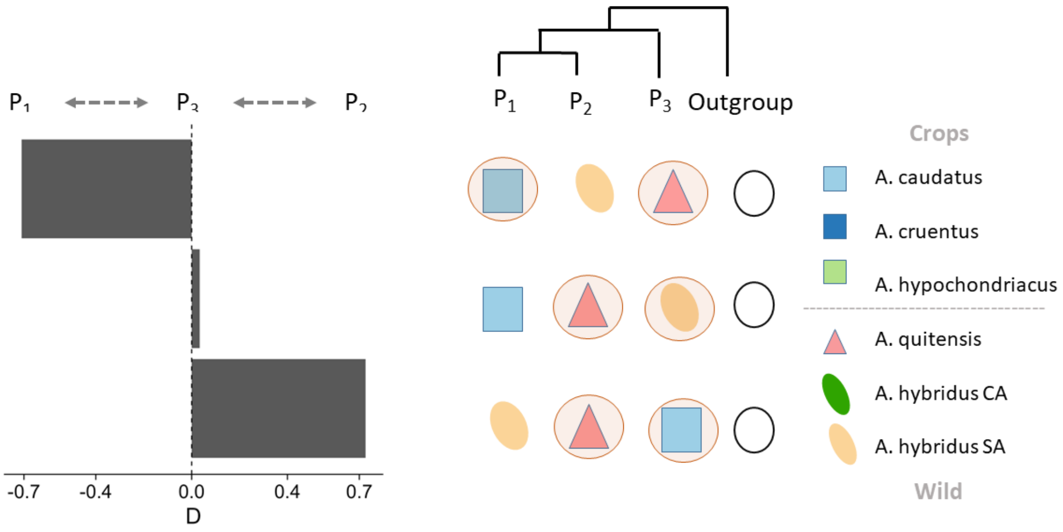


Figure S4: D-statistic value for comparisons between populations of South America. Each bar represents the D-value for each population. The arrows represent the direction of gene flow pairs. Exchanging populations are highlighted with circles. Only significant trees are represented. *A. tuberculatus* was used as an outgroup.

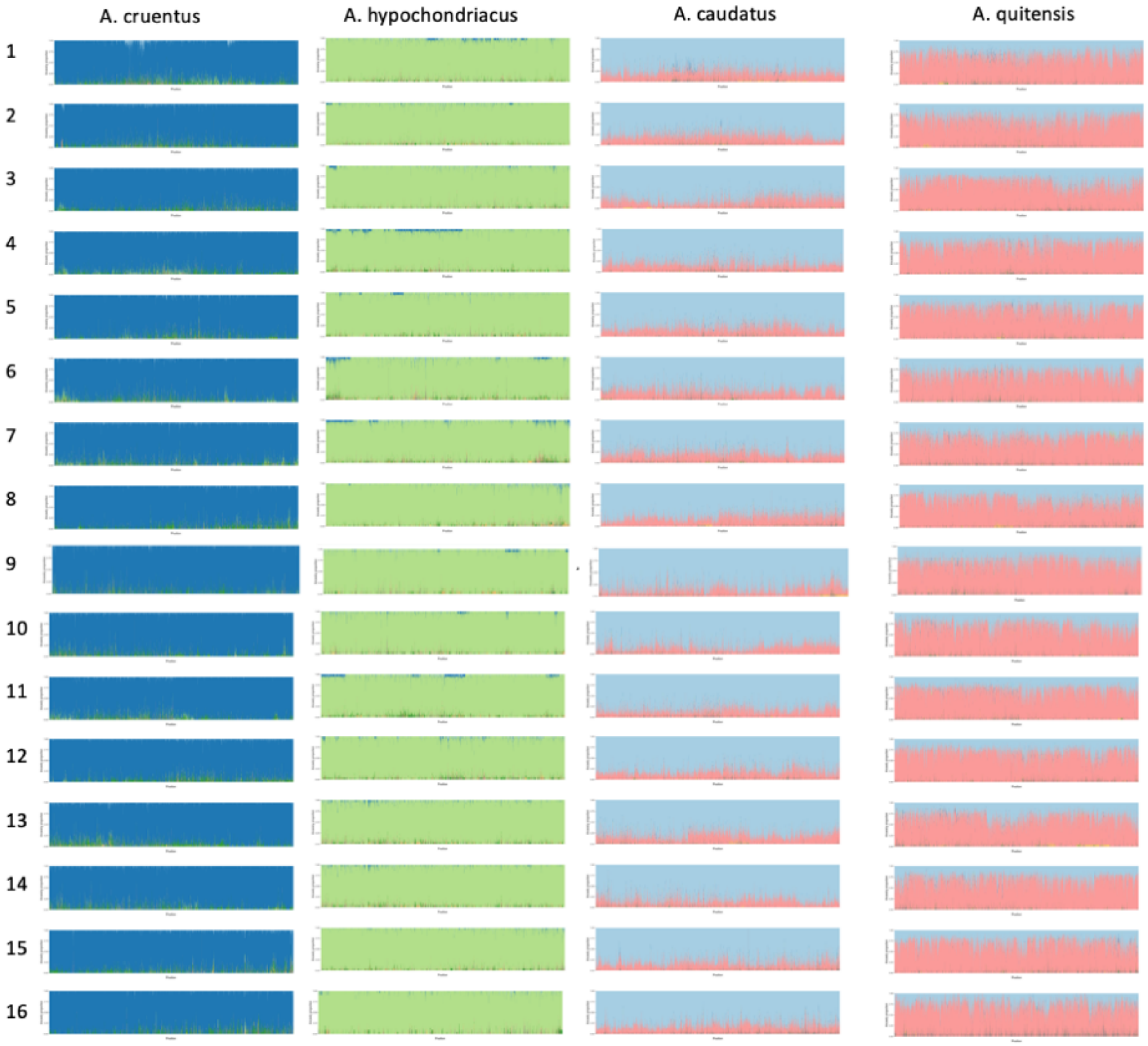


Figure S5: Ancestry proportions summary along the genome, per recipient population, per scaffold. Colors according to Figure 2.

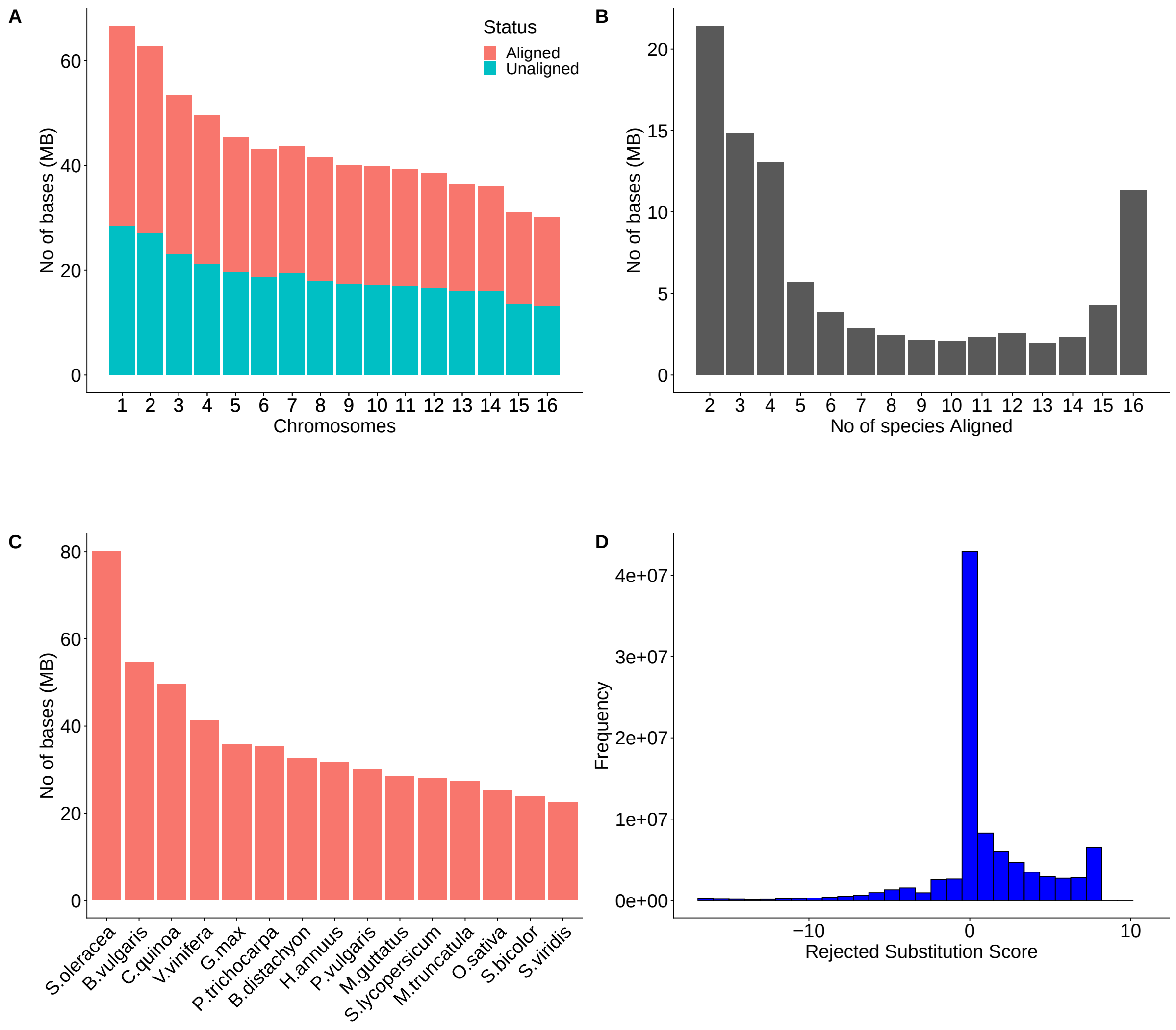


Figure S6: Summary for multiple genome alignment and GERP score. (A) No of bases covered by at least one taxon per Scaffold; (B) Per base coverage of the reference genome by aligning taxon (species); (C) No of bases of the reference genome (*A. hypochondriacus*) covered by each aligning species; (D) Distribution of GERP score (calculated as rejected substitution score).

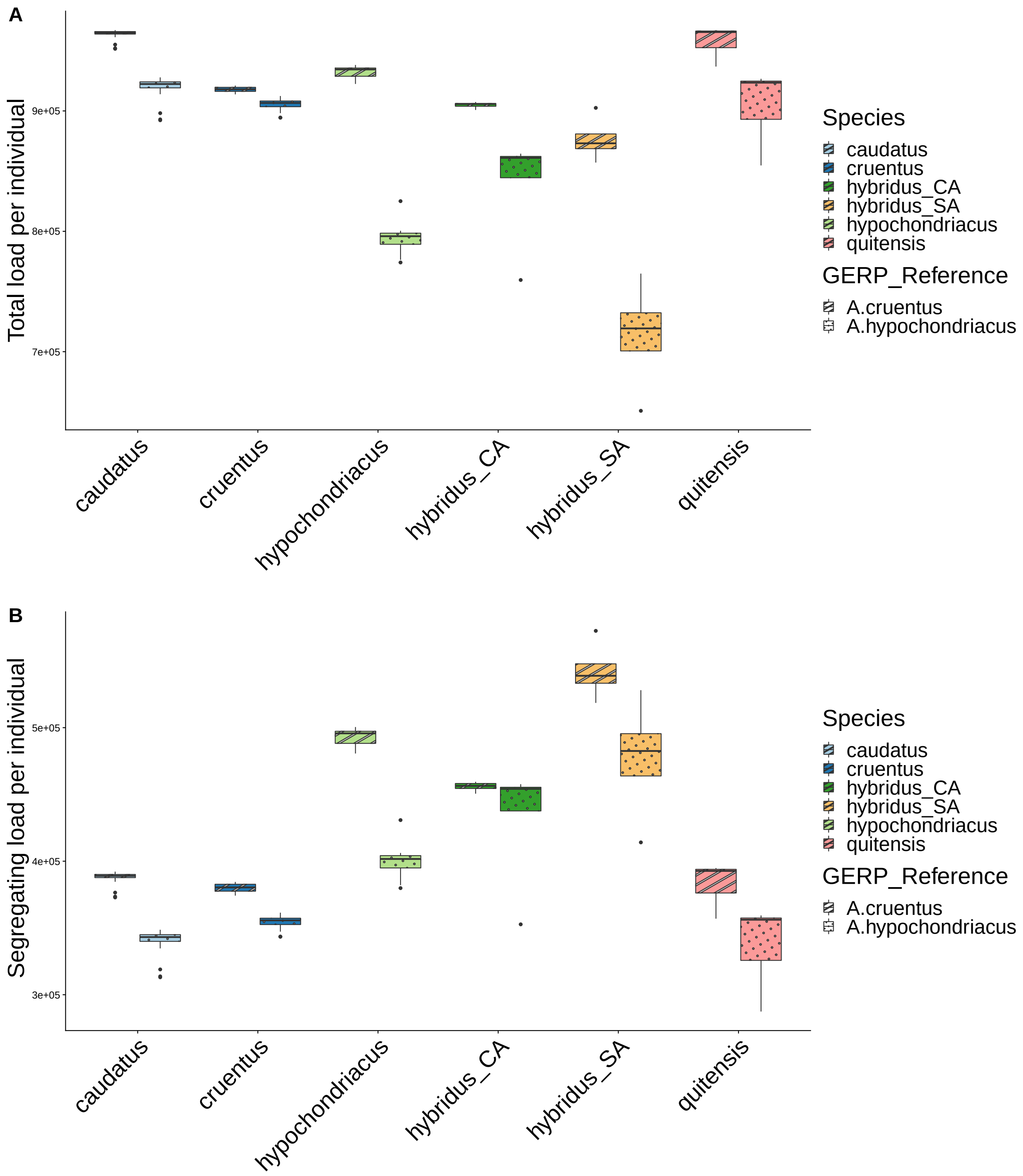


Figure S7: Distribution of genetic load during domestication of grain amaranth using two different genomes for accounting for reference bias. The genetic load was calculated as the sum of GERP scores for derived allele per individual using *A. hypochondriacus* as reference or *A. caudatus* as reference. Distribution of total genetic load per individual in domesticated and wild populations of grain amaranth at (A) all sites (B) segregating sites is depicted using two different reference genomes. The observation of decreased load in hypochondriacus as compared to wild species when using *A. hypochondriacus* genome is not observed when *A. cruentus* is used as a reference for calling GERP. However, the overall pattern for load is comparable for both genomes, suggesting for negligible reference bias.

#### Supplementary Tables

Table S1: **Summary of non-redundant trees to which we found significant D-values.** Note: ANGSD D-Statistic values were updated to positive, and P1 and P2 positions were switched when negative values were present. This was due to make the comparisons easier with other tools as they require comparisons only between populations on P2 and P3.

| P1 | P2 | P3 (target) | D (ANGSD) | D (Dsuite) | D (admixtools) | f4-ratio | f3 (treemx) | f3(admixR) |
|---|---|---|---|---|---|---|---|---|
| hybridus_SA | quitensis | hypochondriacus | 0.225016 | NA | NA | NA | NA | NA |
| caudatus | quitensis | hybridus_SA | 0.033597 | 0.0158596 | 0.0004726645 | 0.00997218 | 0.0887609 | NA |
| caudatus | quitensis | hypochondriacus | 0.037009 | 0.0248318 | 0.0006999955 | 0.00776116 | 0.0831595 | NA |
| caudatus | quitensis | cruentus | 0.032232 | 0.0166697 | 0.0004753609 | 0.0037502 | 0.0755036 | NA |
| cruentus | hybridus_CA | caudatus | 0.228626 | 0.192587 | 0.0097571493 | 0.108917 | 0.0723928 | -0.0068594 |
| hybridus_CA | hypochondriacus | caudatus | 0.10076 | 0.195984 | 0.0140320691 | 0.175783 | 0.0621802 | NA |
| cruentus | hypochondriacus | caudatus | 0.228272 | 0.286137 | 0.0237892185 | 0.265554 | 0.0578926 | NA |
| hybridus_SA | caudatus | hypochondriacus | 0.212658 | NA | NA | NA | 0.028558 | NA |
| hybridus_SA | quitensis | cruentus | 0.17963 | NA | NA | NA | 0.0255764 | NA |
| hybridus_SA | caudatus | cruentus | 0.1691 | NA | NA | NA | 0.0250975 | NA |
| hybridus_SA | quitensis | caudatus | 0.728471 | NA | 0.0894367462 | NA | 0.00870881 | NA |
| hybridus_SA | caudatus | quitensis | 0.70984 | NA | 0.0889640817 | NA | 0.00827429 | NA |

Table S2: Demographic parameters estimated for different demographic scenarios using Fastsimcoal2.

| 3*Parameters | 3*Population Split No gene Flow | 3*Population Split- One time Gene Flow | Population Split Continuous Gene Flow | | |
|---|---|---|---|---|---|
| | | | 2*Point Estimates | 95 % Confidence Interval | |
| | | | | Lower Limit | Upper Limit |
| ANCSIZE1 | 1571884 | 1713678 | 1650048 | 103347 | 2408236 |
| ANCSIZE2 | 1564050 | 446635 | 496443 | 240294 | 2152774 |
| ANCSIZE3 | 152842 | 396708 | 384858 | 10335 | 503711 |
| NHCA | 15444 | 17835 | 20319 | 11200 | 27808 |
| NHSA | 16529 | 16548 | 9637 | 8736 | 22814 |
| NHYP | 14762 | 8349 | 10595 | 5838 | 13133 |
| NCAU | 15146 | 12949 | 8501 | 7099 | 14691 |
| THYP | 8728 | 9894 | 11955 | 5916 | 16802 |
| TCAU | 9553 | 9633 | 5506 | 5065 | 12818 |
| TDIV | 31036 | 30952 | 32832 | 30405 | 40153 |
| TMIGSTART | - | 9157 | - | - | - |
| TMIGEND | - | 9344 | - | - | - |
| MIG23 | - | 3.38801E-05 | 3.19288E-05 | 2.4E-05 | 4.7E-05 |
| MIG32 | - | 3.45086E-06 | 5.39545E-06 | 2E-06 | 9E-06 |
| MaxEstLhood | -83545622.13 | -82852846.15 | -82822883.13 | - | - |
| deltaL | 8294205.304 | 7601429.324 | 7571466.3 | - | - |
| AIC | 384741828.2 | 381551484.9 | 381413496.1 | - | - |
| deltaAIC | 3328332.113 | 137988.8248 | 0 | - | - |

ANCSIZE1 represents ancestral population size of hypochondriacus and hybridus_CA.
ANCSIZE2 represents ancestral population size of caudatus and hybridus_SA.
ANCSIZE3 represents Ancestral population size.
NHCA represents the population size of hybridus_CA.
NHSA represents the population size of hybridus_SA.
NHYP represents the population size of hypochondriacus.
NCAU represents the population size of caudatus.
THYP represents time of split for hypochondriacus
TCAU represents time of split for caudatus
TDIV represents divergence time between hybridus_CA and hybridus_SA
TMIGSTART represents the start time of the migration
TMIGEND represents the end time of the first migration
MIG23 represents migration rate from hypochondriacus to caudatus
MIG32 represents migration rate from caudatus to hypochondriacus
MaxEstLhood maximum log-likelihood of the best estimate.
deltaL difference between estimated and observed log-likelihood.
AIC Akaike's information criterion, AIC = 2d - 2ln(Lhood), where d is the number of parameters.
deltaAIC AIC - min(AIC).

Table S3: Crosses within and between species along with primer pairs used to validate the crosses. The ”intra” represents crosses of plants from the same species (different accessions); ”inter” represents crosses between plants of different species.

| crossing type | maternal species | paternal species | maternal accession | paternal accession | survivale rate (%) | primer pair |
|---|---|---|---|---|---|---|
| inter | A. caudatus | A. cruentus | PI 490518 | PI 511714 | 0 | Primer5 |
| inter | A. caudatus | A. cruentus | PI 490518 | PI 643058 | 0 | Primer5 |
| inter | A. caudatus | A. cruentus | PI 490612 | PI 511714 | 0 | Primer5 |
| inter | A. caudatus | A. cruentus | PI 490518 | PI 643058 | 0 | Primer5 |
| inter | A. caudatus | A. cruentus | PI 490518 | PI 511717 | 0 | Primer5 |
| inter | A. caudatus | A. cruentus | PI 490612 | PI 643058 | 0 | Primer6 |
| inter | A. caudatus | A. hypochondriacus | PI 490518 | PI 558499 | 100 | Primer5 |
| inter | A. caudatus | A. hypochondriacus | PI 490518 | PI 643070 | 100 | Primer5 |
| inter | A. caudatus | A. hypochondriacus | PI 490612 | PI 643070 | 100 | Primer6 |
| inter | A. caudatus | A. hypochondriacus | PI 490612 | PI 558499 | 100 | Primer6 |
| inter | A. caudatus | A. hypochondriacus | PI 490518 | PI 558499 | 100 | Primer5 |
| inter | A. hypochondriacus | A. cruentus | PI 643070 | PI 511714 | 100 | Primer3 |
| inter | A. hypochondriacus | A. cruentus | PI 558499 | PI 643058 | 100 | Primer3 |
| inter | A. hypochondriacus | A. cruentus | PI 643070 | PI 643058 | 0 | Primer3 |
| intra | A. caudatus | A. caudatus | PI 642741 | PI 490518 | 100 | Primer5 |
| intra | A. caudatus | A. caudatus | PI 490612 | PI 490518 | 100 | Primer5 |
| intra | A. caudatus | A. caudatus | PI 490612 | PI 642741 | 100 | Primer6 |
| intra | A. cruentus | A. cruentus | PI 643058 | PI 511714 | 100 | Primer3 |
| intra | A. cruentus | A. cruentus | PI 511714 | PI 643058 | 100 | Primer4 |
| intra | A. cruentus | A. cruentus | PI 511717 | PI 511714 | 100 | Primer2 |
| intra | A. cruentus | A. cruentus | PI 511717 | PI 643058 | 100 | Primer4 |
| intra | A. hypochondriacus | A. hypochondriacus | PI 558499 | PI 604581 | 100 | Primer1 |
| intra | A. hypochondriacus | A. hypochondriacus | PI 604581 | PI 558499 | 100 | Primer1 |
| intra | A. hypochondriacus | A. hypochondriacus | PI 643070 | PI 558499 | 100 | Primer3 |
| intra | A. hypochondriacus | A. hypochondriacus | PI 604581 | PI 643070 | 100 | Primer1 |

Table S4: List of primers used to validate the crosses for the F1 plants

| primer pair | forward primer | reverse primer |
|---|---|---|
| Primer1 | TCACCAATCCCTCCCTCCAA | ACGCGGCGGTTATATGTGAT |
| Primer2 | ACAATTCACATGCAAGCCGG | CCCGTTGCACGATTTTCCAA |
| Primer3 | GACTTGCCTCCTGGAATGCA | AAATCGGTGCAACGTTCTGC |
| Primer4 | GTGACGACAATGATGCTGCC | CGTAACGCATGTGGCATCTG |
| Primer5 | AGTAGACAAACTGGAACCCGA | TGGTCACTTCCAAGGTATGCA |
| Primer6 | AGCTTGTTCAATGCATGGGT | ACGCAACTCTTACAGAGGTCG |

Table S5: List of accessions used in this the study. Names follow USDA germplasm ID

| Name | Country_Origin | Population | ENA_ID |
|---|---|---|---|
| PI 490689 | Ecuador | caudatus | ERR3021332 |
| PI 490739 | Ecuador | caudatus | ERR3021337 |
| PI 490511 | Peru | caudatus | ERR3021351 |
| PI 490491 | Argentina | caudatus | ERR3021382 |
| PI 481949 | Peru | caudatus | ERR3021403 |
| PI 490459 | Bolivia | caudatus | ERR3021366 |
| PI 481957 | Peru | caudatus | ERR3021380 |
| PI 511706 | Peru | caudatus | ERR3021381 |
| AMA 125 | Peru | caudatus | ERR3021385 |
| PI 490561 | Peru | caudatus | ERR3021387 |
| PI 642741 | Bolivia | caudatus | ERR3021388 |
| PI 511704 | Peru | caudatus | ERR3021389 |
| PI 490518 | Peru | caudatus | ERR3021390 |
| PI 649227 | Peru | caudatus | ERR3021394 |
| PI 481960 | Peru | caudatus | ERR3021395 |
| PI 511687 | Peru | caudatus | ERR3021397 |
| PI 490612 | Peru | caudatus | ERR3021398 |
| PI 649217 | Peru | caudatus | ERR3021399 |
| PI 511712 | Peru | caudatus | ERR3021400 |
| PI 490431 | Peru | caudatus | ERR3021401 |
| PI 490609 | Ecuador | caudatus | ERR3021402 |
| PI 511696 | Peru | caudatus | ERR3021404 |
| PI 511686 | Peru | caudatus | ERR3021405 |
| PI 481965 | Peru | caudatus | ERR3021406 |
| PI 511690 | Peru | caudatus | ERR3021407 |
| PI 686455 | Peru | caudatus | ERR3021408 |
| PI 490604 | Bolivia | caudatus | ERR3021409 |
| PI 511679 | Argentina | caudatus | ERR3021430 |
| PI 511680 | Argentina | caudatus | ERR3021431 |
| PI 511681 | Bolivia | caudatus | ERR3021432 |
| PI 608019 | Ecuador | caudatus | ERR3021440 |
| PI 649228 | Peru | caudatus | ERR3021443 |
| PI 649230 | Peru | caudatus | ERR3021444 |
| PI 667165 | Brazil | cruentus | ERR3021328 |
| PI 649509 | Mexico | cruentus | ERR3021329 |
| PI 643042 | Mexico | cruentus | ERR3021422 |
| PI 643039 | Mexico | cruentus | ERR3021410 |
| PI 643058 | Mexico | cruentus | ERR3021411 |
| PI 511713 | Peru | cruentus | ERR3021412 |
| PI 511717 | Guatemala | cruentus | ERR3021413 |
| PI 649514 | Mexico | cruentus | ERR3021414 |
| PI 606798 | Mexico | cruentus | ERR3021415 |
| PI 576482 | Mexico | cruentus | ERR3021416 |

| | | | |
|---|---|---|---|
| PI 649609 | Mexico | cruentus | ERR3021417 |
| PI 451826 | Guatemala | cruentus | ERR3021419 |
| Ames5552 | Mexico | cruentus | ERR3021421 |
| PI 643049 | Mexico | cruentus | ERR3021423 |
| PI 649524 | Mexico | cruentus | ERR3021424 |
| PI 433228 | Guatemala | cruentus | ERR3021429 |
| PI 511714 | Peru | cruentus | ERR3021433 |
| PI 576481 | Mexico | cruentus | ERR3021436 |
| PI 643037 | Mexico | cruentus | ERR3021442 |
| PI 658728 | Mexico | cruentus | ERR3021447 |
| PI 667160 | Guatemala | cruentus | ERR3021448 |
| PI 667158 | Guatemala | hybridus_CA | ERR3021331 |
| PI 511724 | Mexico | hybridus_CA | ERR3021336 |
| PI 604582 | Mexico | hybridus_CA | ERR3021340 |
| PI 604574 | Mexico | hybridus_CA | ERR3021341 |
| PI 604568 | Mexico | hybridus_CA | ERR3021437 |
| PI 490489 | Peru | hybridus_SA | ERR3021333 |
| PI 511754 | Ecuador | hybridus_SA | ERR3021391 |
| PI 686451 | Peru | hybridus_SA | ERR3021426 |
| PI 636180 | Colombia | hybridus_SA | ERR3021441 |
| PI 649537 | Mexico | hypochondriacus | ERR3021343 |
| PI 643036 | Mexico | hypochondriacus | ERR3021344 |
| PI 649575 | Mexico | hypochondriacus | ERR3021345 |
| Ames5457 | Mexico | hypochondriacus | ERR3021346 |
| PI 633589 | Mexico | hypochondriacus | ERR3021347 |
| PI 649565 | Mexico | hypochondriacus | ERR3021348 |
| PI 604559 | Mexico | hypochondriacus | ERR3021349 |
| PI 643070 | Mexico | hypochondriacus | ERR3021350 |
| PI 604581 | Mexico | hypochondriacus | ERR3021352 |
| Ames2085 | Mexico | hypochondriacus | ERR3021353 |
| PI 643041 | Mexico | hypochondriacus | ERR3021354 |
| PI 649602 | Mexico | hypochondriacus | ERR3021355 |
| PI 604587 | Mexico | hypochondriacus | ERR3021356 |
| PI 649607 | Mexico | hypochondriacus | ERR3021357 |
| PI 511731 | Mexico | hypochondriacus | ERR3021359 |
| PI 643067 | Mexico | hypochondriacus | ERR3021360 |
| PI 649559 | Mexico | hypochondriacus | ERR3021361 |
| PI 649595 | Mexico | hypochondriacus | ERR3021362 |
| PI 649623 | Mexico | hypochondriacus | ERR3021364 |
| PI 604595 | Mexico | hypochondriacus | ERR3021439 |
| PI 649529 | Mexico | hypochondriacus | ERR3021445 |
| PI 511749 | Ecuador | quitensis | ERR3021365 |
| PI 511737 | Ecuador | quitensis | ERR3021367 |
| PI 490664 | Ecuador | quitensis | ERR3021335 |
| PI 667156 | Ecuador | quitensis | ERR3021338 |
| PI 490705 | Ecuador | quitensis | ERR3021370 |
| PI 490684 | Ecuador | quitensis | ERR3021342 |

| | | | |
|---|---|---|---|
| PI 691596 | Argentina | quitensis | ERR3021368 |
| PI 649246 | Peru | quitensis | ERR3021369 |
| PI 511745 | Ecuador | quitensis | ERR3021374 |
| PI 652426 | Brazil | quitensis | ERR3021371 |
| PI 490720 | Ecuador | quitensis | ERR3021376 |
| PI 511741 | Ecuador | quitensis | ERR3021377 |
| PI 669830 | Peru | quitensis | ERR3021378 |
| PI 511751 | Peru | quitensis | ERR3021372 |
| PI 652428 | Brazil | quitensis | ERR3021373 |
| PI 652422 | Brazil | quitensis | ERR3021375 |
| PI 490466 | Peru | quitensis | ERR3021392 |
| PI 490673 | Ecuador | quitensis | ERR3021393 |
| PI 669836 | Argentina | quitensis | ERR3021379 |
| PI 490670 | Ecuador | quitensis | ERR3021334 |
| PI 490679 | Ecuador | quitensis | ERR3021396 |
| PI 669839 | Peru | quitensis | ERR3021428 |
| PI 511736 | Bolivia | quitensis | ERR3021434 |
| PI 511747 | Ecuador | quitensis | ERR3021435 |
| ERR3220318 | | tuberculatus | ERR3220318 |
